# Epigenetic histone modifications H3K36me3 and H4K5/8/12/16ac induce open polynucleosome conformations via different mechanisms

**DOI:** 10.1101/2024.02.19.580980

**Authors:** Yi-Yun Lin, Peter Müller, Evdoxia Karagianni, Willem Vanderlinden, Jan Lipfert

**Affiliations:** Department of Physics and Center for NanoScience (CeNS), LMU Munich, Amaliensstrasse 54, 80799 Munich, Germany; Soft Condensed Matter and Biophysics, Department of Physics and Debye Institute for Nanomaterials Science, Utrecht University, Princetonplein 1, 3584 CC Utrecht, The Netherlands; School of Physics and Astronomy, University of Edinburgh, James Clerk Maxwell Building, Peter Guthrie Tait Road, Edinburgh EH9 3FD, United Kingdom

**Keywords:** Nucleosomes, chromatin, methylation, acetylation, single-molecule methods

## Abstract

Nucleosomes are the basic compaction unit of chromatin and nucleosome structure, and their higher-order assemblies regulate genome accessibility. Many post-translational modifications alter nucleosome dynamics, nucleosome-nucleosome interactions, and ultimately chromatin structure and gene expression. Here, we investigate the role of two post-translational modifications associated with actively transcribed regions, H3K36me3 and H4K5/8/12/16ac, in the contexts of tri-nucleosome arrays that provide a tractable model system for quantitative single-molecule analysis, while enabling us to probe nucleosome-nucleosome interactions. Direct visualization by AFM imaging reveals that H3K36me3 and H4K5/8/12/16ac nucleosomes adopt much more open and loose conformations than unmodified nucleosomes. Similarly, magnetic tweezers force spectroscopy shows a reduction in DNA outer turn wrapping and nucleosome-nucleosome interactions for the modified nucleosomes. The results suggest that for H3K36me3 the increased breathing and outer DNA turn unwrapping seen in mononucleosomes propagates to more open conformations in nucleosome arrays. In contrast, the even more open structures of H4K5/8/12/16ac nucleosome arrays do not appear to derive from the dynamics of the constituent mononucleosomes, but are driven by reduced nucleosome-nucleosome interactions, suggesting that stacking interaction can overrule DNA breathing of individual nucleosomes. We anticipate that our methodology will be broadly applicable to reveal the influence of other post-translational modifications and action of nucleosome remodelers.

## Introduction

Nucleosomes are the basic building block of eukaryotic genomes, essential for the organization, compaction, and regulation of genetic information [1–3]. Canonical nucleosome core particles are composed of two copies of H2A, H2B, H3, and H4 assembled into a histone octamer that is wrapped by 147 bp of DNA [2, 4] (Figure 1A). Interaction within nucleosomes stems from both electrostatic interactions and specific molecular contacts [5–9]. The nucleosome core interacts with adjacent nucleosomes to form the higher order structure, so that, ultimately, the genomic DNA on a scale of ∼1 m can be packed and condensed into the nucleus, which is on a scale of ∼µm [10–16]. However, the DNA must remain accessible for various cellular processes such as replication, transcription, and repair [17–21]. Multiple factors affect the compaction and chromatin structure to regulate those cellular processes. Epigenetic modifications, or post-translational modification (PTMs), a diverse array of covalent chemical marks that modulate gene expression without altering the DNA sequence, have emerged as critical regulators of chromatin architecture and function [22–26]. In eukaryotic cells, histones are subject to hundreds of PTMs including acetylation, methylation, ubiquitination, phosphorylation, and sumoylation [27]. Histones PTMs are widely distributed throughout the whole genome. They can control the accessibility of DNA or recruit chromatin remodelers to regulate gene expression [22–26, 28–30]. Histone PTMs are present both in the tails of histones and their globular core domains [31, 32]. By introducing additional charge, neutralizing existing charge, or by adding steric constraints, different modifications affect the compaction of chromatin and also modulate the stability of nucleosomes. In particular, methylation and acetylation have been intensively studied as marks of chromatin status involving active or silenced transcription [25, 27]. For acetylation (“ac”), histone acetylation neutralizes the positive charge of lysine, which reduces interactions with DNA and has been shown to e.g., enable transcription factor binding within nucleosomes [33–35]. Acetylation of H4 tail has a strong effect on weakening chromatin packing *in vivo* and *in vitro* [33, 36–38]. H3 acetylation also reduces the charge of the tails but the effect on folding propensity of nucleosome arrays is less clear [35, 39].

**Figure 1.**
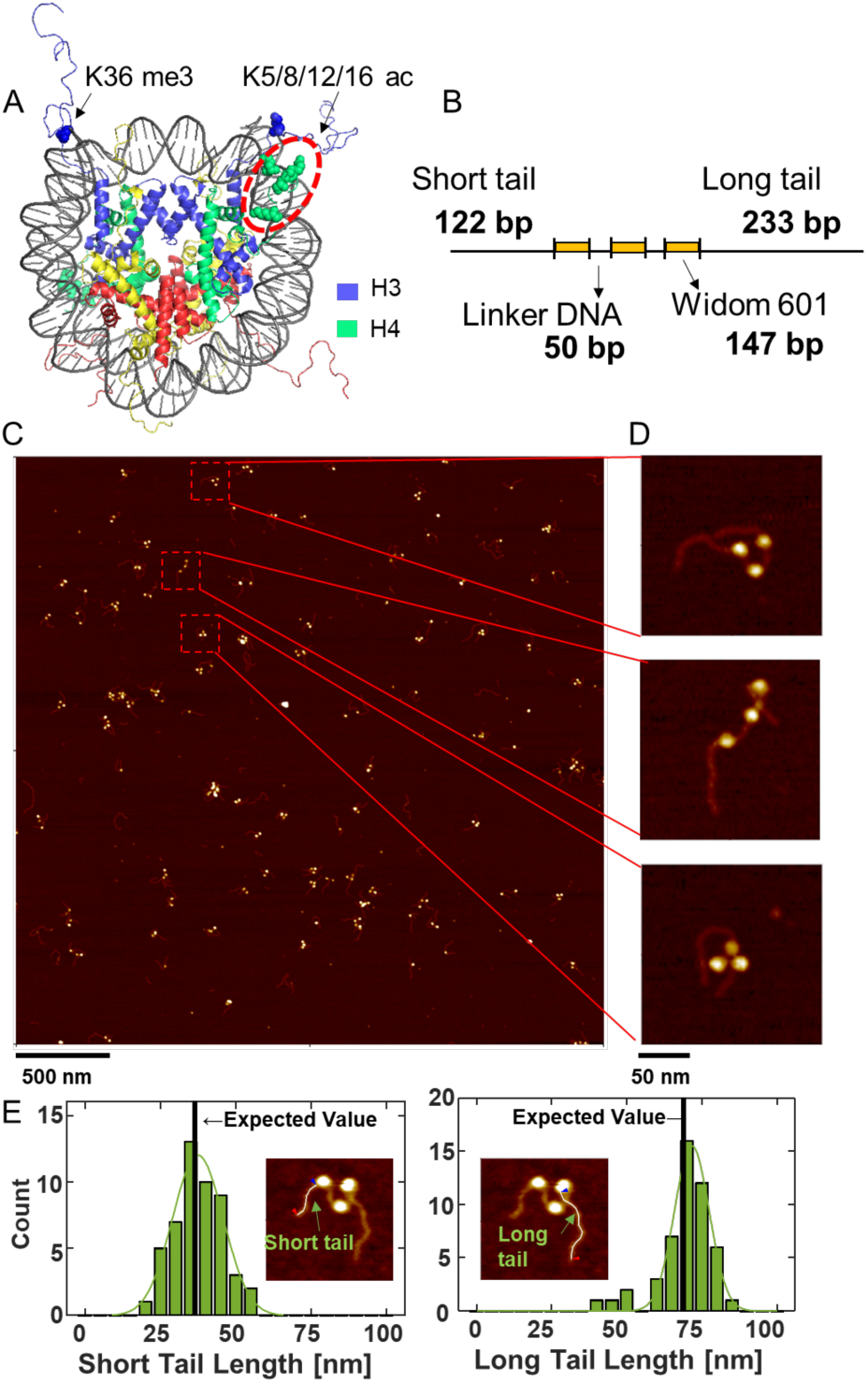
Analysis of tri-nucleosome conformations by AFM imaging. **A)** Crystal structure of a canonical nucleosome (PDB 1KX5). Colored spheres represent the positions of the modified amino acids in the histone tail. Residues involve in H3K36me3 (i.e. three additional methyl groups at lysine 36 of histone H3) shown as a blue sphere and in H4K5/8/12/16 ac (i.e. acetylation of H4 histones at lysines 5, 8, 12 and 16) as green spheres) **B)** Schematic of the DNA construct used for AFM imaging. The 896 bp DNA consists of three 147 bp Widom 601 nucleosome positioning sequences that are flanked by a short and a long arm of 122 bp and 233 bp, respectively. **C)** AFM image of DNA and tri-nucleosome sample with a field of view of 3 µm x 3 µm (recorded with 2048 x 2048 pixels). **D)** Zooms of selected tri-nucleosomes in the AFM image in panel C. **E)** Histograms of short and long arm length of unmodified tri-nucleosomes, with a Gaussian fitted to each distribution (green solid line). Insets show example image of tri-nucleosomes with the poly-line profile indicated that was used to measure the arm lengths. Vertical lines are the expected arm length computed from the number of base pairs in the short and long arm, respectively, and assuming 0.314 ± 0.13 nm/bp.

Histone methylation (“me”) occurs on the side chains of lysines or arginines [40] and, unlike acetylation, does not alter the charge of histone protein and is thought to act mainly via “reader” enzymes that specifically recognize the methylated site and then activate or repress transcription [41]. For example, H3K9 and H3K27 methylation are often related to silenced chromatin states [41]. Examples of chromatin readers that recognize methylation and are involved in gene repression are HP1 that binds to H3K9me3 and contributes to heterochromatin formation [42, 43] and the methyltransferase PRC2 that acts on H3K27 [44] and recruits other accessory protein to propagate the H3K27me3 mark resulting in gene silencing [45–47]. In contrast, H3K36 methylation is associated with actively transcribed regions [48, 49].

While traditional biochemical and structural methods have provided valuable insights into nucleosome architecture, these approaches often entail ensemble measurements that obscure the intrinsic heterogeneity and dynamic nature of these macromolecular assemblies. Recently, single-molecule techniques have provided an ability to probe nucleosomes at the level of individual molecules [50–54]. In particular, AFM imaging has been used to visualize the structure and dynamics of nucleosomes and their interactions [55–63]. We have recently developed a high-throughput pipeline to image individual nucleosomes [57, 59] and applied the approach to determine the effect of several epigenetic modifications on mononucleosome conformational landscape. AFM imaging of mononucleosomes revealed that H3K36me3 nucleosomes are, on average, more open and wrap less DNA, while H3S10 phosphorylation and H4K5/8/12/16ac did not significantly affect conformations of individual nucleosomes [58]. A complementary approach has been to probe nucleosomes and nucleosome arrays by force spectroscopy, in optical [64–67] and magnetic tweezers [52, 68], which enable to apply forces and monitor the resulting changes in extension [52, 66, 68, 69]. Force-spectroscopy approaches have revealed changes in extension in intermediate nucleosome conformations and characterized the folding of chromatin fibers and higher order assemblies [52, 66–68, 70, 71].

Here, we go beyond mononucleosomes and probe the effect of epigenetic modifications on nucleosome-nucleosome interactions using arrays with three nucleosomes, where the conformational landscape of individual nucleosomes is modulated and constrained by interactions. We complement our AFM imaging results using magnetic tweezers force-spectroscopy measurements [72–77]. By applying mechanical forces and observing the ensuing responses, we can probe nucleosome conformations and interactions and go beyond the static structures revealed by AFM imaging [55–59, 78]. Our single-molecule results consistently indicate that both H3K36me3 and H4K5/8/12/16ac lead to more open conformations in the context of tri-nucleosome constructs, by reducing stacking interactions and increasing nucleosome breathing.

## Results and Discussion

### Assembly and AFM imaging of tri-nucleosome arrays

To prepare nucleosome samples for AFM imaging, we assembled different variant nucleosomes by salt gradient dialysis on 895 bp DNA constructs. Our DNA construct features three Widom 601 (W601) sequences [79] partitioned by 50 bp of linker DNA and flanked by a short arm 120 bp and long arm 232 bp (Figure 1B). The same DNA construct was used for the different nucleosome variants. We deposited nucleosome samples on poly-L-lysine coated mica and recorded high-resolution AFM images (see Materials and Methods for details). AFM images (Figure 1C) are obtained by amplitude modulation AFM in air and further analyzed to dissect the influences of PTMs on structural dynamics and geometry. The AFM images show populations of naked DNA, mono-, di-, and tri-nucleosomes (Supplementary Figure 1). We designed the DNA construct with two different length arms flanking the region with the W601 sequences to be able to determine nucleosome positioning. To quantify the positioning, we first evaluate the length of the two arms for individual tri-nucleosome particles (see Materials and Methods for details) (Figure 1D,E). The length of the short arm and long arm are 37.3 ± 8.4 nm and 71.6 ± 6.4 nm, respectively. These results are in an excellent agreement with the expected values of 38 nm and 73 nm for short and long arm, assuming a DNA length per base pair of 3.14 ± 0.13 Å found previously by AFM imaging under similar conditions [57], fully consistent with positioning of the nucleosomes on the W601 sequences.

We use AFM imaging to confirm the assembly of different variant nucleosomes and quantify the different polynucleosome populations, by counting the number of mono-, di-, tri-, and even occasional tetra-nucleosomes (requiring nucleosome loading to DNA outside of the W601 sequences) that are successfully assembled (Supplementary Figure 1). The populations for bare DNA, and DNA with one, two, three, and four nucleosomes are consistent, within experimental errors, with a simple binomial distribution (Supplementary Figure 1), which implies that the assembly of the different variant nucleosomes on the three W601 sites are all relatively uncooperative under the conditions of our experiments, consistent with previous observations [78, 80]. We find similar probabilities *P* for sites being occupied for the different variants, with nearly identical values for unmodified (*P* = 0.418 ± 0.010) and H4K5/8/12/16ac (*P* = 0.415 ± 0.008). H3K36me3 exhibits a slightly lower occupation probability of *P* = 0.344 ± 0.008, which might be due to minor differences in the protein concentration due to experimental variability or due to a slightly lower affinity of the tri-methylated variant. Overall, AFM imaging confirms that nucleosomes of all three variants are assembled robustly on our DNA construct, with similar affinities and relatively low cooperativity between positing sites.

### AFM imaging reveals conformational changes of tri-nucleosome arrays induced by epigenetic modifications

To study the effect of selected PTMs on nucleosome structure, we analyze the configuration of tri-nucleosomes by extracting several structure parameters from AFM images. In a first step, we use process images in SPIP and identify the tri-nucleosome samples (Supplementary Figure 2). Nucleosome positions are ordered from the nucleosome closest to the short tail to the one closest to long tail (referred to as N_1_, N_2_, N_3_) and we extract the x and y positions of the nucleosome centers (Supplementary Figure 2C). As a first geometric parameter to quantify tri-nucleosome conformations, we calculate the distance between the first nucleosome and the third, which we call the N_1_N_3_-distance. In addition, we compute the inner angle α, defined as the angle between the lines connecting N_2_ to N_1_ and N_3_, using the formula 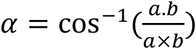 with *a* and b are two dimensional vectors of the nucleosome particles (Supplementary Figure 2D, 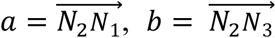). Finally, we determine the radius of gyration defined by 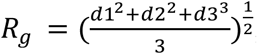 where (*d*1*, d*2*, d*3) are the distances from the nucleosome positions to their center of mass (Supplementary Figure 2C).

We apply AFM imaging and image analysis to obtain distributions of the geometric parameters to quantify and compare the impact of the different PTMs (Figure 2). For each modification, we measure the radii of gyration as a parameter describing the overall nucleosome distances in the tri-nucleosome complex (Figure 2A,D,G), the distance between the outer two nucleosomes (Figure B,E,H), and the angle at the inner nucleosome (Figure 2C,E,I). To facilitate direct comparison of the impact of the nucleosome types, we smooth histograms for a given parameter using a kernel density estimate (Figure 2A-I) and co-plot the resulting probability densities (Figure 2J-L).

**Figure 2.**
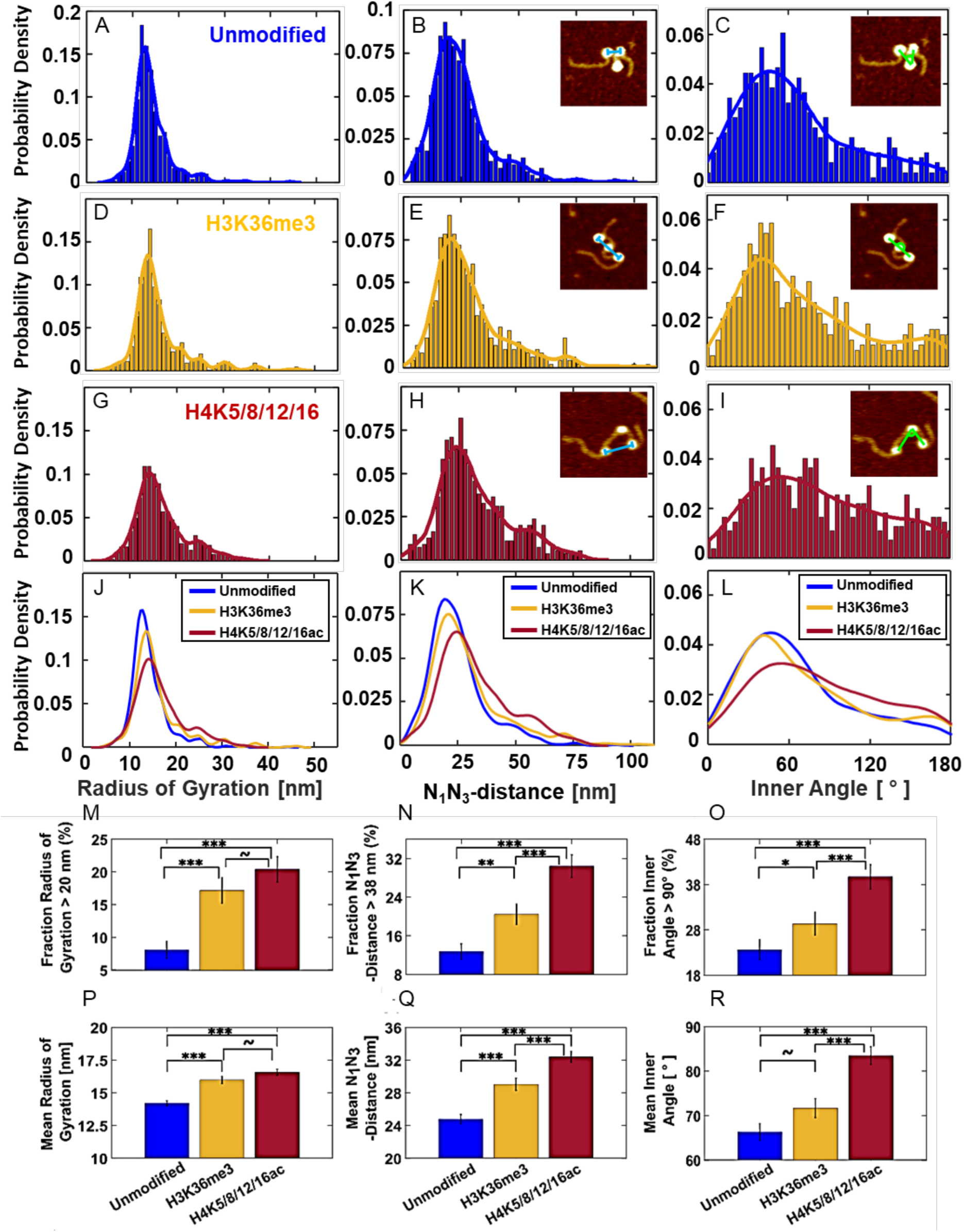
AFM imaging reveals the impact of H3K36me3 and H4K5/8/12/16ac PTMs on tri-nucleosome conformations. **A-I)** Probability distributions for radii of gyration, N_1_N_3_-distances, and inner angles determined from AFM imaging for unmodified (A-C), H3K36me3 (D-F), and H4K5/8/12/16ac (G-I) nucleosomes. Raw data are shown in the histogram. Solids lines are kernel density estimated. **J-L)** show the kernel density estimates of the different nucleosome modifications co-plotted for each parameter for ease of comparison. The numbers of molecules analyzed for each condition are indicated in panels A, D, and G. The insets in panels B, C, E, F, H, and I show example AFM images of tri-nucleosome constructs with the N_1_N_3_-distances (blue line) and inner angles (green line) indicated. **M-R)** Quantitative comparisons of the distributions from panels (A-L). Panels M-O show the fraction of particles exhibiting values larger than a given threshold for the different parameters as indicated in the y-axis labels for the different nucleosome types, i.e. the fraction of tri-nucleosomes adopting a more “open” configuration. Error bars indicate the counting error. Bars between the columns indicate the results of a two-tail two-sample proportion-test. (P-R) Mean values of the parameter distribution for the different nucleosome types. The bars between the columns indicate the result of a two-tail two-sample t-test. Error bars indicate the standard error of the mean. ∼ not significant, *p < 0.05, **p<0.01, ***p<0.001.

We find that the radii of gyration, N_1_N_3_-distances, and inner angles provide a highly consistent picture: The unmodified nucleosomes exhibit the most compact conformations, exhibiting narrow distributions, with the smallest mean values for all three parameters. Conversely, H4K5/8/12/16ac nucleosomes present the broadest range and largest average values, while H3K36me3 nucleosomes exhibit distributions for radii of gyration, N_1_N_3_- distances, and inner angles that are intermediate between the other two nucleosome types (Table 1 and Figure 2A-L). Together, these data suggest that H4K5/8/12/16ac nucleosomes exhibit the least compact and most open conformations, while unmodified nucleosomes exhibit the most compact structures and H3K36me3 nucleosomes take on intermediate conformations. Comparing the mean values for radii of gyration, N_1_N_3_-distances, and inner angles, we find statistically significant differences (assessed by two-sample *t*-tests) with unmodified nucleosomes being most compact and H4K5/8/12/16ac taking on the largest values, except for the radii of gyrations comparison between H3K36me3 and H4K5/8/12/16ac and the inner angle comparison between unmodified and K3K36me3, which are not significant (Figure 2P-R). In addition to comparing the overall distributions and their means, we look at the subpopulations with open conformations, defined as having *R*_g_, N_1_-N_3_- distance, or inner angle α values above a manually determined threshold (Figure 3M-O). The fraction of particularly open conformations increases, in almost all cases statistically significantly, in going from unmodified, to H3K36me3, and further to H4K5/8/12/16ac nucleosomes, further confirming the observations from the overall distributions (Figure 3M-O). In addition to comparing the means, we also compare the full distribution using Kolmogorov–Smirnov tests to compare the full distribution of *R*_g_, N_1_-N_3_-distance, and inner angle α between the different variant nucleosomes (Supplementary Table 1). We find statistically significant differences for all parameters (N_1_N_3_-distance, *R*_g_, and inner angle α) and for each pairwise comparison, except for the inner angle comparison between unmodified and H3K36me3.

**Figure 3.**
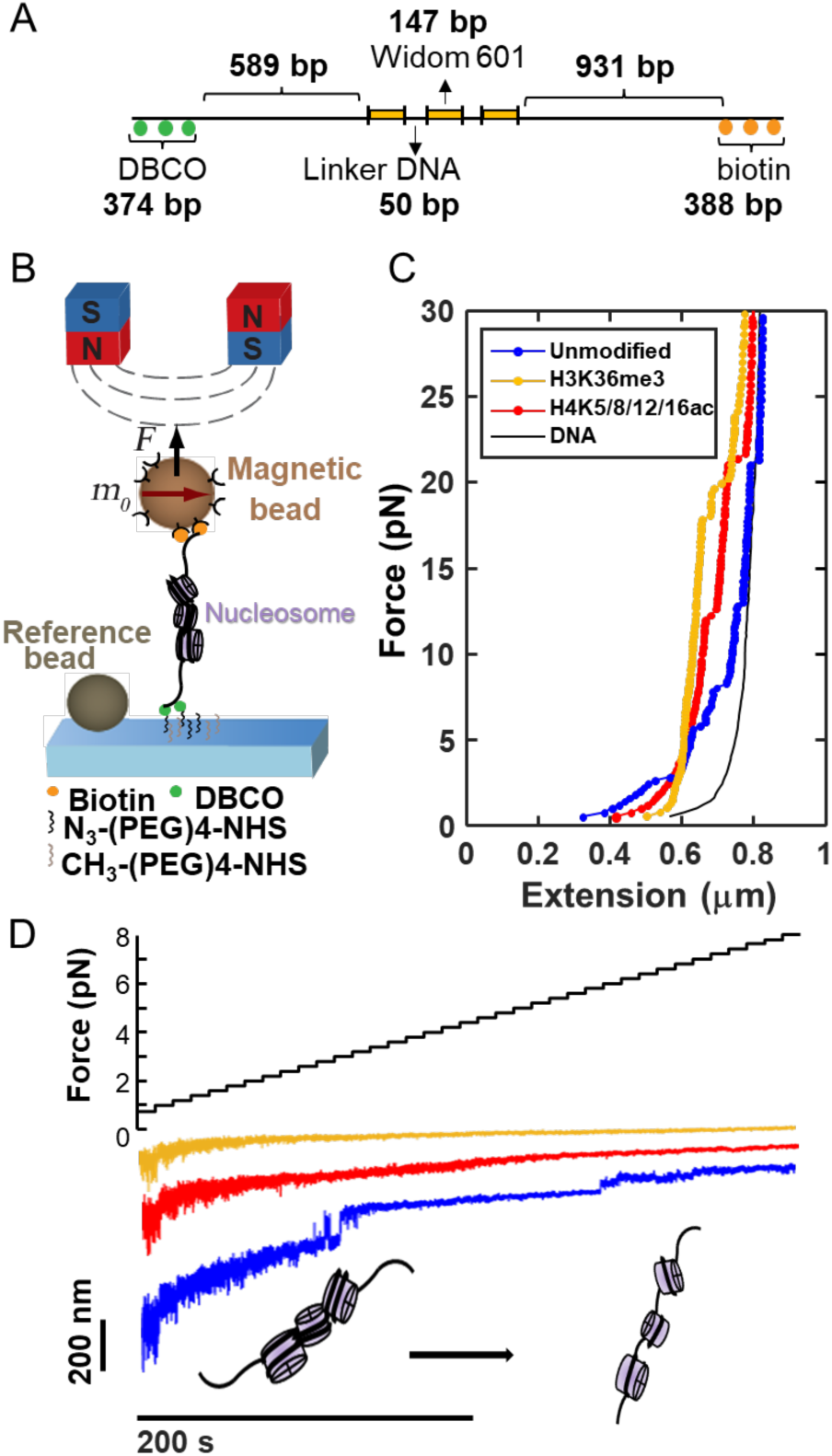
Probing tri-nucleosomes in magnetic tweezers. **A)** Schematic of the DNA construct used for magnetic tweezers. The 2823 bp DNA consists of three 147 bp Widom 601 nucleosome positioning sequences that are flanked by a 589 bp short arm and extra 374 bp fragment with DBCO labeled, and a 931 bp long arm and extra 388 bp fragment with biotin labeled, respectively. **B)** Schematic of the magnetic tweezer set-up. Nucleosomes are reconstituted on DNA with two functionalized ends, one labeled with multiple biotins and the other with multiple DBCOs. The DNA construct is amplified by using the ligation free “megaprimer” method described previously [80]. The flow cell surface is functionalized with azide-(PEG)_4_-NHS. The magnetic beads are labeled with streptavidin. **C)** Force-extension curves of different variant nucleosomes and bare DNA anchored as shown in panel A. Nucleosome samples were stretched under applied forces from 0.5 to 30 pN. **D)** Force ramp at low force (Force ≤ 8 pN; top) of different variants of nucleosome. The extension time traces (color curves; bottom) show different length plateaus at forces ≤ 8 pN that indicate outer turn unwrapping and unstacking of polynucleosomes. Same color code as in panel C.

**Table 1:** Mean value and standard deviation (SD) of three parameter: Radius of gyration (Rg), N_1_N_3_-Distance (N_1_N_3_), and inner angle (Angle) from Figure 2.

|  | Unmodified | H3K36me3 | H4K5/8/12/16ac |
| --- | --- | --- | --- |
| **Number of tri-nucleosomes** | 495 | 460 | 549 |
| ***Mean Rg (nm)*** | 14.2 | 16.0 | 16.6 |
| ***SD Rg (nm)*** | 4.2 | 6.0 | 5.4 |
| ***Mean N<sub>1</sub>N<sub>3</sub> (nm)*** | 24.8 | 29.0 | 32.4 |
| ***SD N<sub>1</sub>N<sub>3</sub> (nm)*** | 12.6 | 16.3 | 15.3 |
| ***Mean Angle (°)*** | 66.3 | 71.7 | 83.5 |
| ***SD Angle (°)*** | 41.8 | 46.0 | 46.5 |

The more open configurations for H3K36me3 compared to unmodified tri-nucleosome arrays are in line with the behavior of the constituent mononucleosomes. Previous work using a high-throughout AFM analysis approach to probe mononucleosomes found that H3K36me3 mononucleosomes have increased breathing activity, are almost 2-fold less likely to occupy the fully wrapped state and exhibit less anti-cooperativity for unwrapping from the respective ends compared to unmodified nucleosomes [58]. In contrast, the same assay found no difference between the conformations of H4K5/8/12/16ac and unmodified mononucleosomes, in stark contrast to our findings for tri-nucleosomes.

To be able to even more directly compare how mononucleosome conformations vary across the different PTMs under the conditions of our assay, we exploit the fact that in our tri-nucleosome samples there is a sub-population of molecules with only one nucleosome assembled (Supplementary Figure 1). We analyzed this sub-population of mononucleosomes by tracing the DNA entry/exit angles (Supplementary Figure 3). From the analysis of the mononucleosome sub-population in our tri-nucleosome data, we find that H3K36me3 nucleosomes have statistically significant larger mean exit angles compared to unmodified and H4K5/8/12/16ac nucleosomes, while there is no significant difference between the unmodified and H4K5/8/12/16ac condition, in excellent agreement with the previous analysis using mononucleome samples assembled on shorter DNA with only one W601 positioning sequence [57].

Together, the observations suggest that the acetylation of H4K5/8/12/16ac nucleosomes primarily affects nucleosome-nucleosome interactions and the open, more dynamic conformations of H4K5/8/12/16ac tri-nucleosome mostly occur due to a decrease in stacking and/or binding interactions between the nucleosomes, compared to unmodified and H3K36me3. Our experimental observations for H4K5/8/12/16ac tri-nucleosomes are in good agreement with molecular simulations that investigated histone tail acetylation dependence of the free energy landscape of tri-nucleosome and found that tri-nucleosomes with H4 acetylation have a larger *R*_g_ compared to unmodified nucleosomes and also reduce the contact between first and third nucleosomes mediated by the histone tails [81]. Our results support that H4-acetylation opens nucleosome array by reducing the inter-nucleosome interaction [82]

### Effect of the ion atmosphere on tri-nucleosome conformations

Since chromatin structure is sensitive to the ionic environment [83–88], we performed control AFM imaging measurement using a different buffer composition and compared the structural parameter in the presence of different types of salt. It is well-known that Mg^2+^ can affect the compaction of chromatin [83, 86, 89, 90]. Mg^2+^ can help chromatin to turn from ‘beads-on-a-string’ into a 30 nm fiber *in vitro* [89] and Mg^2+^ and K^+^ mixed environment seems important for the structure of heterochromatin formation [91]. Previous work by sedimentation velocity analytical ultracentrifugation on nucleosome arrays in different mixed salt solution shows that the additions of Mg^2+^ leads to the precipitation of nucleosome arrays in solution with KCl or NaCl [85]. Therefore, we compared the effect of the mixed ionic Mg^2+^ and K^+^ (2 mM MgCl_2_ and 100 mM KCl; which is approximately the physiological concentration of ions intracellularly) and used as the buffer for the measurements described above, to the 200 mM NaCl buffer condition, the standard deposition buffer employed previously to characterize the effect of PTMs on single nucleosomes [58]. The results show that both unmodified tri-nucleosome and the acetylated tri-nucleosome adopt more compact structures in the presence of Mg^2+^ and K^+^ (Supplemtary Figure 4), in line with previously observed trends for chromatin. However, the effect of the change in ionic conditions is smaller than the effect of the PTMs on structure. In fact, the change induced by changing from the NaCl imaging buffer to the mixed conditions with Mg^2+^ was smaller, for all parameters analyzed, than the difference between unmodified and H4K5/8/12/16ac nucleosomes (Supplementary Figure 4). In conclusion, while we find that the addition of Mg^2+^ compacts tri-nucleosome arrays in agreement with previous findings, the observed influence of PTMs on the structure of tri-nucleosome is similar for different salt conditions and dominates under the conditions employed here.

**Figure 4.**
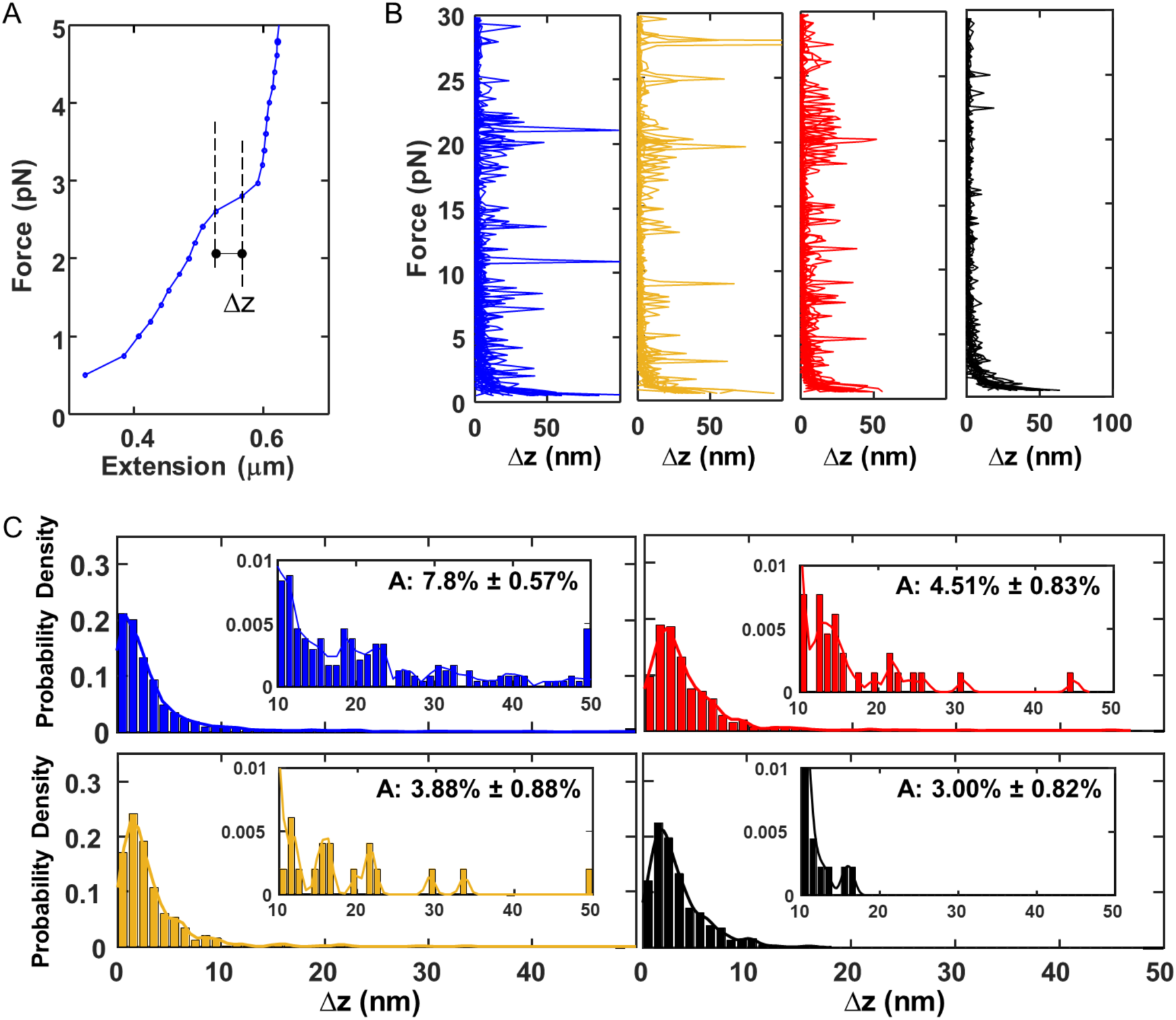
Analysis of force-extension reveals nucleosome unstacking and unwrapping. **A)** Schematic of the Δz analysis, using the unmodified nucleosome force-extension curve from Figure 3B. We analyzed the force-extension data for polynucleosomes by averaging each force plateau’s z positions and subtracting the average z position from the previous force plateau to obtain Δz. **B)** Δz vs. force data for unmodified (*N*=19), H3K36me3 (*N*=16), H4K5/8/12/16 (*N*=21), and DNA (*N*=15). **C)** Histograms of Δz values with kernel density estimates (solid lines) using the data in the force range 2-8 pN for different variant nucleosomes and bare DNA. The insets show histograms of Δz from 10 – 50 nm with kernel density estimates and the fractions of counts for Δz ≥ 10 nm. A indicates the area under the curve for Δz from 10-50 nm. The indicated error is the counting error. The fraction of events with Δz ≥ 10 nm is significantly lower for H3K36me3 or H4K5/8/12/16ac compared to unmodified nucleosomes (*p* = 0.00036 and *p* = 0.0016, respectively), while the difference between the two PTMs is not significant (*p* = 0.45), based on two-sample two-tailed proportion tests.

### Magnetic tweezers force spectroscopy probes unmodified, H4K5/8/12/16ac, and H3K36me3 tri-nucleosome constructs

Having established that H3K36me3 and H4K5/8/12/16ac influence internucleosome interactions and result in more open polynucleosome structures, we asked how the PTMs affect the properties of the tri-nucleosomes in a dynamic setting by using multiplexed magnetic tweezers. To study the behavior of variant nucleosomes under controlled stretching forces, we assembled nucleosomes on a 2823 bp DNA construct with biotin labels on one and DBCO labels at the other end, separated by unmodified DNA from a central segment containing 3x W601 and 50 bp of linker DNA each (Figure 3A). To produce sufficient DNA for *in vitro* nucleosome reconstitution and with appropriate labels for stable attachment in the tweezers, we used our megaprimer approach describe previously [80] and reconstituted unmodified, H3K36me3, and H4K5/8/12/16 tri-nucleosomes on the construct. The biotin labels enable attachment to streptavidin-coated magnetic beads (M270, 2.7 µm diameter), while the DBCO labeled-end provides covalent attachment to an azide-functionalized glass slide surface via copper-free click chemistry [80] (Figure 3A). To confirm the assembly of nucleosomes and to quantify the different polynucleosome populations, we again used AFM imaging to count the number of successfully assembled mono-, di-, tri-nucleosomes (Supplementary Figure 5). The distributions show similar binomial distributions as we observed for the assembly on shorter length DNA used for AFM analysis discussed above.

We performed force-extension experiments on polynucleosome arrays by applying constant forces in the magnetic tweezers from 0.5 to 30 pN in 0.2 pN increments, each for 5 s (for forces > 8 pN) or 10 s (≤ 8 pN). The raw extension traces reveal considerable variability for all variant polynucleosomes (Figure 3C and Supplementary Figure 6), showing the heterogeneity and complexity of our reconstituted samples, in line with our AFM imaging results. The time traces also reveal that, superimposed on the expected force-extension stretching response of double-stranded DNA, there are jumps and hopping events visible in the data, qualitatively in line with nucleosomes unwrapping and unstacking. We compare the different tri-nucleosome constructs by taking the mean extension for each force plateau to obtain force-extension curves (Figure 3B). We find that at low forces (≤ 8 pN), the unmodified nucleosome tethers tend to have a shorter extension compared to H3K36me3 and H4K5/8/12/16ac. In addition, the raw extension vs. time traces below for forces ≤ 8 pN show that unmodified nucleosomes exhibit larger fluctuations due to hopping or stepping contributions compared to H3K36me3 and H4K5/8/12/16ac constructs (Figure 3C and Supplementary Figure 6). At high forces (> 8 pN) all types of nucleosomes show steps with comparable properties.

### Repeated stretching and release cycles indicate that mechanical forces disrupt some but not all nucleosome interactions

We observe clear differences in the tether responses between the first stretching cycle (going from 0.5 to 30 pN) and the first release or second stretching cycle (Supplementary Figure 7). After the first stretching cycle, the tether lengths at a given force are increased compared to the initial stretching cycle for all nucleosome variants investigated, suggesting that at least some of the nucleosome structures are permanently disrupted by applying forces, in agreement with previous literature [64, 92]. Nonetheless, repeated force-extension cycles still show steps and a decreased extension, compared to bare DNA, at low forces, implying that some nucleosomes remain bound or can rebind even after stretching to 30 pN, consistent with previous observations that the core particle may reassemble upon relaxation after peeling off the inner turn DNA [53, 64, 66, 93, 94].

### Force spectroscopy suggests a reduction of stacking and outer turn wrapping interactions in H4K5/8/12/16ac and H3K36me3 compared to unmodified nucleosomes

The time traces in the low force regime (≤ 8 pN) exhibit a broad range of steps, hopping behavior, and gradual changes in extension, while the traces at higher forces show more clearly defined steps. We attribute the changes in the range of 2-8 pN to unwrapping of the outer turn of DNA from nucleosomes and the disruption of nucleosome-nucleosome interactions. In contrast, the defined steps at high forces (> 8 pN) agree with previous reports [52, 53, 68, 69, 92, 95–101] of non-equilibrium peeling of the inner ∼75 bp of DNA from the core of the octamer. Here, we first discuss the behavior at low forces (≤ 8 pN) and in the next section we analyze the steps at higher forces.

To compare the different variant nucleosomes in force-extension measurements, we compute the mean extension in z for each force plateau and calculate the difference in z between adjacent force steps (Figure 4A), which we define as Δz. Spikes in Δz correspond to abrupt jumps in tether lengths (Figure 4B). The computed Δz values show that all types of nucleosomes demonstrate multiple spikes from low to high forces (Figure 4B). Unmodified nucleosomes have higher density of spikes, and the spikes are distributed over a broader range of forces. For both H3K36me3 and H4K5/8/12/16ac, the spikes are less dense at low force regime (≤ 8 pN) compared to the unmodified condition. We analyze the Δz distribution at forces ranging from 2 to 8 pN. The result shows that H3K36me3 and H4K5/8/12/16ac have narrower distributions and a reduced population of events with Δz ≥ 10 nm compared to unmodified nucleosomes (Figure 4C). By calculating the relative population for Δz ≥ 10 nm, we find that H3K36me3 (3.88% ± 0.88%) and H4K5/8/12/16ac (4.51% ± 0.83%) exhibit significantly fewer large steps than unmodified nucleosomes (7.80% ± 0.57%).

The reduced number of stepping events in the force range 2-8 pN for H3K36me3 and H4K5/8/12/16ac compared to unmodified tri-nucleosomes suggests that these PTMs disrupt nucleosome-nucleosome stacking and outer turn wrapping. The magnetic tweezers observations are in line with AFM results that indicate more open and diverse and open conformations for H3K36me3 and H4K5/8/12/16ac. Interestingly, while the AFM results suggest that H4K5/8/12/16ac tri-nucleosomes adopt the most open conformations, the magnetic tweezers measurements see the smallest proportion of steps for H3K36me3. However, we note that the difference in Δz steps > 10 nm between H3K36me3 and H4K5/8/12/16ac is within experimental error.

### Force spectroscopy finds no influence of the investigated PTMs on inner turn unwrapping of nucleosomes

The Δz vs. force plots from variant nucleosome also show that at higher forces (> 8 pN), there are multiple spikes regardless of nucleosome types (Figure 4B). The corresponding steps are consistent with inner turn nucleosome unwrapping. To quantify the effects of the investigated PTMs on inner turn unwrapping, we analyzed the extension steps at high forces (> 8 pN) with the step finding algorithm by Kerssemakers *et al.* [102] to identify unwrapping steps in our extension vs. time traces (Figure 5A). From the fits, we determine the differences of average extensions before and after the steps to obtain step sizes. The distributions of step sizes from the three different types of nucleosomes show very similar peaks with mean step sizes between 21-24 nm (Figure 5B), in excellent agreement with previous reports for step sizes of inner turn unmodified nucleosome unwrapping in the range of 20-30 nm [52, 53, 68, 69, 92, 95–101]. In addition, we analyze the forces at which the high-force steps occur to quantify the force range of inner turn unwrapping. We again find remarkably similar force distributions for all types of nucleosomes studied, with mean forces well within experimental error, at 19-20 pN (Figure 5C). The results suggest that the H3K36me3 and H4K5/8/12/16ac PTMs have no significant effects on inner turn nucleosome disassembly. Inner turn nucleosome unwrapping is sudden due to the strong interactions near to positions ±40 bp of DNA from the dyad axis [103]. Overall, the interactions between the inner turn DNA wrap and the histone octamer involve both electrostatic and non-electrostatic interactions, while the outer DNA wrap interactions with the histone octamer are dominated by electrostatic interactions [96, 103]. Consequently, the changes at the N-tail due to H4K5/8/12/16ac or H3K36me3 are unlikely to affect the inner turn nucleosomal DNA unwrapping, consistent with our experimental findings.

**Figure 5.**
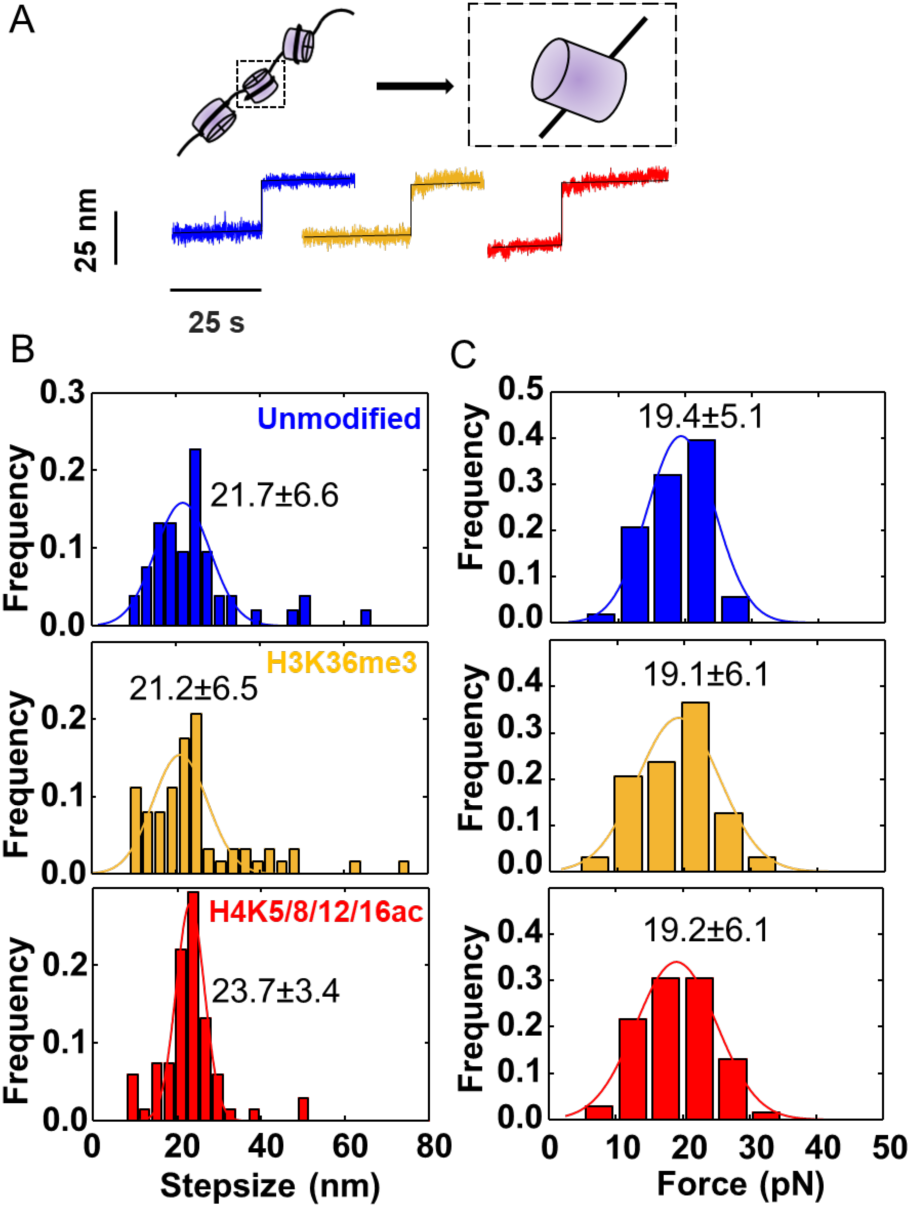
Analysis of inner turn DNA unwrapping in tri-nucleosome constructs under force. **A)** Example of a discrete steps in time traces (colored data) at forces > 8 pN, characteristic of the unwrapping of the inner DNA turn from nucleosomes. Black lines are fitted steps using algorithm by Kerssemakers *et al.* [102]. Unmodified nucleosome, blue line; H3K36me3 nucleosome, yellow line; and H4K5/8/12/16ac nulceosome, red line. **B)** Histograms of the step sizes for inner turn unwrapping as determined in panel A. Solid lines are Gaussian fits and the means are indicated in the panels. **C)** Histogram of the forces for inner turn unwrapping corresponding to the steps in panel B. Solid lines are Gaussian fits and the means are indicated in the panels. The differences between the means in panel B and C for the different nucleosome types are not significant, as assessed by two-tail two-sample *t*-tests.

## Conclusion

PTMs are a key factor that affects the structure and dynamics of chromatin fibers in the cell. They can have manifold effects on chromatin structure, such as entry site unwrapping, nucleosome destabilization, formation of active or repressive compartments, and histone-histone destabilization [31, 32, 104]. Here we investigate the conformations of post-translational modified nucleosomes using two single-molecule techniques: atomic force microscopy imaging and magnetic tweezer force spectroscopy. Specifically, we study the effects of the post-translational modifications H3K36me3 and H4K5/8/12/16ac on tri-nucleosome array structure and mechanical stability. We use tri-nucleosomes, which has been reported to be the smallest cluster size found in cells [105], as a tractable model system for comparison of different PTMs on nucleosome arrays that build in complexity on our previous work on mononucleosomes in isolation [58].

H3K36me and H4K5/8/12/16ac are known as markers of active region in chromatin. Previous high-throughput AFM image analysis has shown that H3K36me3 mononucleosomes exhibit partial unwrapping and more open conformations compared to unmodified mononucleosomes, likely due to the position of the H3K36me3 mark at the DNA entry/exit site of the nucleosome [29, 92]. We confirmed this finding by analyzing the mononucleosome sub-population of our tri-nucleosome samples and find significantly higher exit angles for H3K36me3 nucleosomes compared to unmodified and H4K5/8/12/16ac species. It has been found that PTMs at the entry/exit region enhance partial DNA unwrapping [31, 58, 106]. Our results here suggest that the increased unwrapping induced by the H3K36me seen in mononucleosomes propagates to higher order nucleosome assemblies, as we observe more open and loose conformations for H3K36me3 compared to unmodified nucleosomes both by AFM imaging magnetic and tweezers force spectroscopy. Our findings are in line with previous simulations that predict nucleosome breathing to affect their higher order structures, to result in more heterogeneous nucleosome-nucleosome contacts [107].

The H3K36me3 modification is associated with DNA repair, alternative splicing, and transcription. It is enriched in the region of actively transcribed genes [108–110]. Our finding that H3K36me3 leads to more open nucleosome array structures highlights a mechanism how it can facilitate access of histone-binding proteins, e.g. of protein carrying a PWWP domain [111] that interact with the H3K36me3 mark and regulate gene transcription [108, 110].

Interestingly, the H4K5/8/12/16ac modification causes no significant changes in mononucleosome structure compared to unmodified mononucleosome [58], yet it leads to the most open and extended tri-nucleosome structures as judged by the AFM imaging results, of the three variants studied. This is consistent with the view that the H4K5/8/12/16ac mark, which is known to be associated with open chromatin conformations [33, 112], reduces nucleosome-nucleosome interactions and stacking. The fact that H4K5/8/12/16ac tri-nucleosomes are more open and less compact than the H3K36me3 constructs suggests that nucleosome-nucleosome interactions can be more important and overrule nucleosome breathing and outer turn unwrapping.

*In vitro* work reveals that H4K5/8/12/16ac inhibits liquid-liquid phase separation, likely due to the decrease of multivalent interaction with other nucleosomes [113]. Our experiments are consistent with this observation of reduced liquid-liquid phase separation by H4K5/8/12/16ac nucleosomes, as we observe reduced nucleosome-nucleosome. In contrast, the inner turn unwrapping appears to not be influenced by the investigated PTMs, consistent with the view that their influence is limited to the entry-exit site and tail regions.

Chromatin architecture is more open at transcriptionally active sites [114]. Here we demonstrate that epigenetic marks associated with active transcription can decrease chromatin compaction directly – not only by reducing nucleosome-nucleosome interactions but also by outer turn wrapping affinity. Taken together, our work suggests that the combination of force spectroscopy and AFM imaging can provide a comprehensive understanding of how different PTMs affect nucleosome assemblies and we anticipate our approach to be powerful to study the effect of other PTMs in the future.

## Materials and Methods

### DNA preparation

We prepared two different DNA constructs for the AFM and MT measurements, respectively. We used the plasmid pFMP218, a custom-built plasmid provided by Prof. Felix Müller-Planitz, TU Dresden, Germany) as template to produce DNA constructs with 3 repeats of the Widom 601 sequence [79]. The DNA construct for AFM measurements has a length of 896 bp. We prepared the DNA by PCR with Phusion Hot Start polymerase (follow the vendor’s protocol) by using forward primer 5’-TAAGTTGGGTAACGCCAGG-3’ and reverse primer 5’-GGCCGATTCATTAATGCAGC-3’. The functionalized DNA constructs used for MT measurement were prepared as previously described using a “megaprimer” approach [80]. Two functionalized DNA strands with 50% biotin-16-dUTP or 50% DBCO-(PEG)_4_-dUTP replacement for dTTP, respectively, were obtained by PCR amplification. The two functionalized DNAs were used as “megaprimers” to amplify the final 2823 bp DNA construct with DBCO and biotin labels at the two opposite ends, respectively.

### Nucleosome reconstitution

Nucleosomes were assembled on the labeled DNA construct obtained using the megaprimer protocol outlined in the previous section. Unmodified and modified histone proteins were purchased from EpiCypher (Durham, North Carolina). We followed the previously published protocol to prepare nucleosome reconstitutions [80]. Nucleosome reconstitutions were performed by salt gradient dialysis. The preparation of nucleosome samples for AFM measurement and MT measurement used dialysis chambers containing 2.8 – 3 μg of 896 or 2823 bp DNA at 2 M NaCl that were placed in 300 ml high-salt buffer (2 M NaCl, 10 mM Tris, 1 mM EDTA). 3 liters of low-salt buffer (50 mM NaCl, 10 mM Tris, 1 mM EDTA) were transferred to the high-salt buffer at 4 °C overnight.

### AFM sample preparation, imaging, and analysis

We followed the previously published protocol to prepare samples for AFM imaging [57–59, 115, 116]. The reconstituted nucleosomes were incubated in 100 mM KCl, 2 mM MgCl_2_, and 10 mM Tris-HCl, pH 7.6, for 1 min on ice and then deposited on poly-L-lysine (0.01% w/v) coated muscovite mica for 30 s, followed by 20 ml Milli-Q water rinsing and drying with a gentle stream of filtered N_2_ gas. AFM imaging was performed on a Nanowizard Ultraspeed 2 (JPK, Berlin,Germany) with AFM cantilevers 240AC-NA (Opus) in air. All AFM images were acquired in tapping mode at room temperature. The scans were recorded at 1 Hz line frequency over a field of view of 3 µm x 3 µm at 2048 x 2048 pixels. For image processing, Scanning Probe Image Processor (SPIP v6.5.1; Image Metrology) was employed. Image processing involved background correction by using global fitting with a third-order polynomial and line-by-line correction through the histogram alignment routine.

### Magnetic tweezers setup

We used a custom-built MT setup described previously [117]. The setup was equipped with a pair of 5 x 5 x 5 mm^3^ permanent magnets (W-05-N50-G, Supermagnete, Switzerland) with a 1 mm gap in vertical configuration [118]. In the setup, a DC-motor (M-126.PD2, Physik Instrumente, Germany) controls the distance between magnets and the flow cell. A LED (69647, Lumitronix LED Technik GmbH, Germany) was used for illumination. In addition, a 40x oil-immersion objective (UPLFLN 40x, Olympus, Japan) and a CMOS sensor camera with 4096 x 3072 pixels (12M Falcon2, Teledyne Daisa, Canada) were utilized to image a field of view of 400 x 300 µm^2^. Images were recorded at 58 Hz and transferred to a frame grabber (PCIe 1433; National Instruments, Austin TX). By tracking images in real-time with custom-written tracking software (Labview, National Instruments), we can extract the (x,y,z) coordinates of all beads [119]. The objective is mounted on a piezo stage (Pifoc P726. 1CD, PI Physikinstrumente) to build a look-up table (LUT) for tracking the bead z-position. With a step size of 100 nm, the LUT was generated over a range of 10 µm. Set up control and bead tracking used Labview routines described previously [119].

### Flow cell assembly and preparation

Flow cells were assembled from two microscope cover slips with a parafilm spacer. The bottom coverslip (24 x 60 mm, Carl Roth, Germany) was treated with 2% APTES to generate an aminosilanized surface. Before flow cell assembly, 5000x diluted stock solution of polystyrene beads with 1 μm diameter (Polysciences, USA) in ethanol (Carl Roth, Germany) was deposited on the amino-coated coverslip and then slowly dried. These immobile surface-bound beads serve as reference beads for drift correction. The bottom coverslip was aligned with a pre-cut parafilm and a top coverslip with two small holes for inlet and outlet. Then the assembled flow cell was baked at 80 °C for 1 min to create a seal.

### DNA or polynucleosome anchoring for magnetic tweezers experiments

DNA or polynucleosome anchoring was carried out as described [80]. Briefly, following flow cell assembly, 50 mM each of azide-(PEG)_4_-NHS (Jena Biosciences GmbH, Jena, Germany) and methyl-(PEG)_4_-NHS (Life technologies) in 1 x PBS were introduced and incubated for 1 h [120]. We mixed our DNA or polynucleosome sample in measurement buffer MB1 (MB1; 10 mM HEPES pH 7.6, 100 mM KCl, 2 mM MgCl_2_, 0.1% Tween-20). Next, DNA or polynucleosome were dissolved in 100 μl MB1, flushed into the flow cell and incubated for 1 h. Afterwards, we rinsed with MB2 buffer, which consists of MB1 supplemented with 0.1% (w/v) bovine serum albumin (Carl Roth, Germany) to flush away unbound nucleosome or DNA. Subsequently, we flowed in 1% casein for nucleosome samples or 1.5% (w/v) bovine serum albumin for DNA samples in MB2 into the flow cell, incubated for 1 h to minimize nonspecific interactions. Finally, we flushed in streptavidin-coated M270 beads (Dynabeads, Invitrogen) and incubated with samples to form tethers. After flushing away free magnetic beads with several cell volumes of MB2, we start the measurements.

## Author contributions

Yi-Yun Lin: conceptualization, data analysis, investigation (AFM and MT measurements), project administration, visualization, writing – original draft preparation. Peter Müller: data analysis, investigation (AFM measurements), visualization. Evdoxia Karagianni: data analysis, investigation (MT measurements). Willem Vanderlinden: conceptualization, supervision, project administration, visualization, writing – review and editing. Jan Lipfert: conceptualization, funding acquisition, supervision, project administration, writing – review and editing.

## Acknowledgements

We thank Thomas Nicolaus, Dave van den Heuvel, Relinde van Dijk-Moes, and Elleke van Harten for laboratory assistance, Lori van de Cauter, Sebastian Konrad, Tine Brouns, Philipp Korber, and Felix Müller-Planitz for useful discussions, Stefanie D. Pritzl, Ivo Vermaire and Diogo Saraiva for support with measurements, and Friedrich Förster, Louris Feitsma, Joke Granneman, Mariska Gröllers-Mulderij, and the entire Structural Biochemistry group at Utrecht University for useful discussion and laboratory use.

## Funding

This work was supported by the Deutsche Forschungsgemeinschaft (DFG, German Research Foundation) through SFB 863, Project 111166240 A11, and Utrecht University.

## Notes

### Competing Interest Statement

The authors have declared no competing interest.

