## Supplementary Information for "Epigenetic histone modifications H3K36me3 and H4K5/8/12/16ac induce open polynucleosome conformations via different mechanisms"

**Content:**

**Supplementary Table 1**

**Supplementary Figures 1-7**

### SUPPLEMENTARY TABLES

| Type1 | Type2 | Parameter | P-value |
|---|---|---|---|
| **Unmodified** | H3K36me3 | *R*g | 1.2815e-06 |
| **Unmodified** | H3K36me3 | $N_1N_3$-distances | 0.0039 |
| **Unmodified** | H3K36me3 | Inner angle | 0.1624 |
| **Unmodified** | H4K5/8/12/16ac | *R*g | 5.5441e-15 |
| **Unmodified** | H4K5/8/12/16ac | $N_1N_3$-distances | 2.5401e-15 |
| **Unmodified** | H4K5/8/12/16ac | Inner angle | 4.2401e-09 |
| **H3K36me3** | H4K5/8/12/16ac | *R*g | 9.9286e-04 |
| **H3K36me3** | H4K5/8/12/16ac | $N_1N_3$-distances | 5.6997e-06 |
| **H3K36me3** | H4K5/8/12/16ac | Inner angle | 2.0555e-05 |

**Supplementary Table 1:** Pairwise comparisons by a two-sample Kolmogorov–Smirnov test between nucleosome types for radius of gyration (*R*g), $N_1N_3$-distance, and inner angle.

**SUPPLEMNTARY FIGURES**

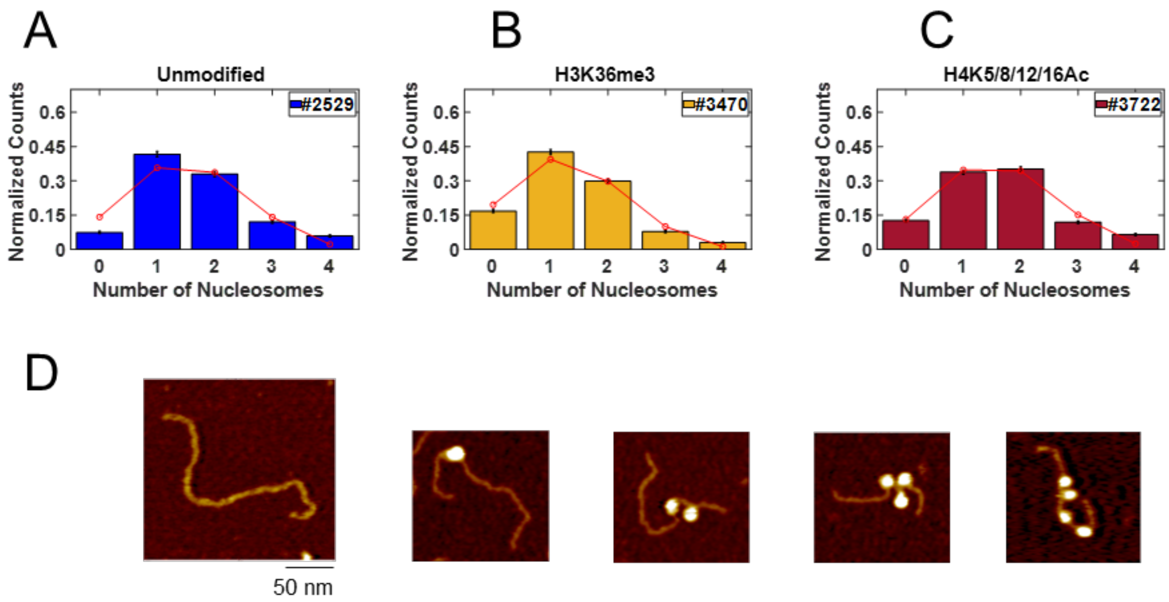


**Supplementary Figure 1. Quantification of nucleosome assembly on DNA from AFM images. A)-C)** Histogram of the number of nucleosomes assembled on DNA strands for (A) unmodified (N = 2529), (B) H3K36me3 (N=3470), and (C) H4K5/8/12/16Ac (N = 3722) obtained from AFM images. The numbers in the legend indicate the number of DNA strands analyzed for each type of nucleosome. Red points are the best fit of a binomial distribution with fitted assembly probability for unmodified (P = 0.418 ± 0.010), H3K36me3 (P = 0.344 ± 0.008), H4K5/8/12/16ac (P = 0.415 ± 0.008). Error bars here indicates the counting error, errors of the fitted P values are the standard error of a binomial distribution. **D)** Examples of AFM images of bare DNA, and unmodified type mono-, di-, tri-, and tetra- nucleosomes. The scale bar applies to all AFM images in panel D.

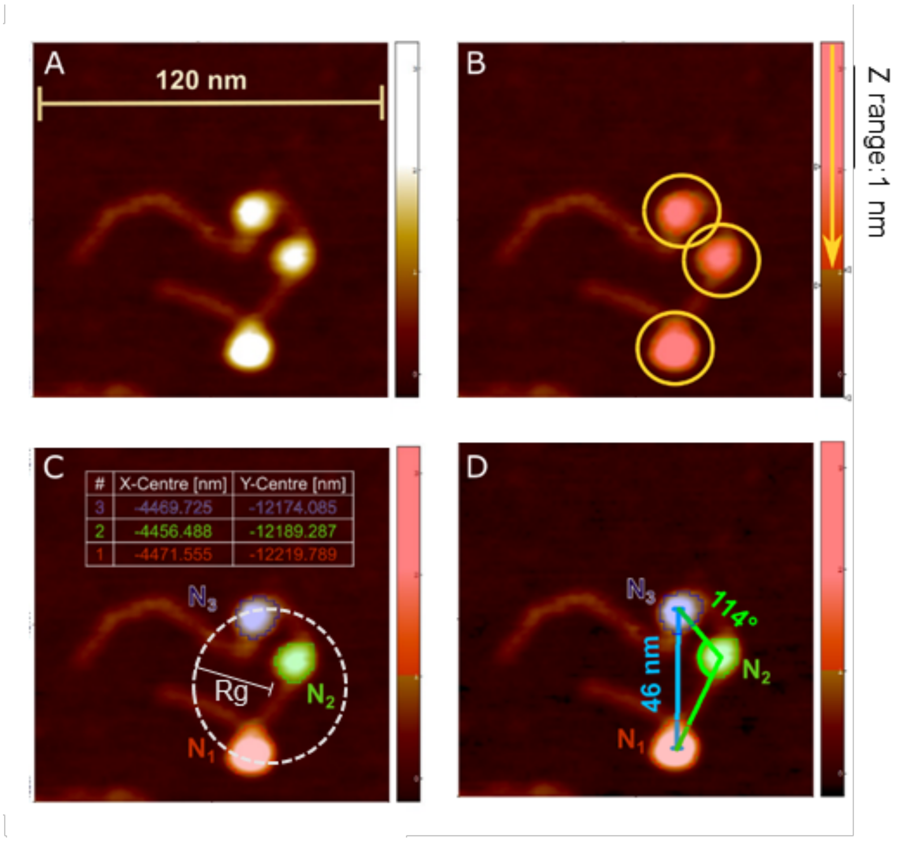


**Supplementary Figure 2. Determination of geometric parameters from AFM images of tri-nucleosomes.** Data analysis was performed in SPIP (Image Metrology). **A)** AFM image of a selected tri-nucleosome sample. **B)** A height threshold (Z range) is adjusted to detect the nucleosomes as areas of the topographic scan where this height limit is exceeded. **C)** Using the particle detection feature the nucleosomes are then identified as three separate particles within the scan and their center positions are displayed in a table. The center positions are used to calculate the radius of gyration ($R_g$). **D)** The nucleosomes are labeled $N_1$, $N_2$, and $N_3$ following the DNA strand from short to long arm. For analysis we determine the $N_1N_3$-distance (blue) and the angle between the outer nucleosome centers and the center of the middle nucleosome (green).

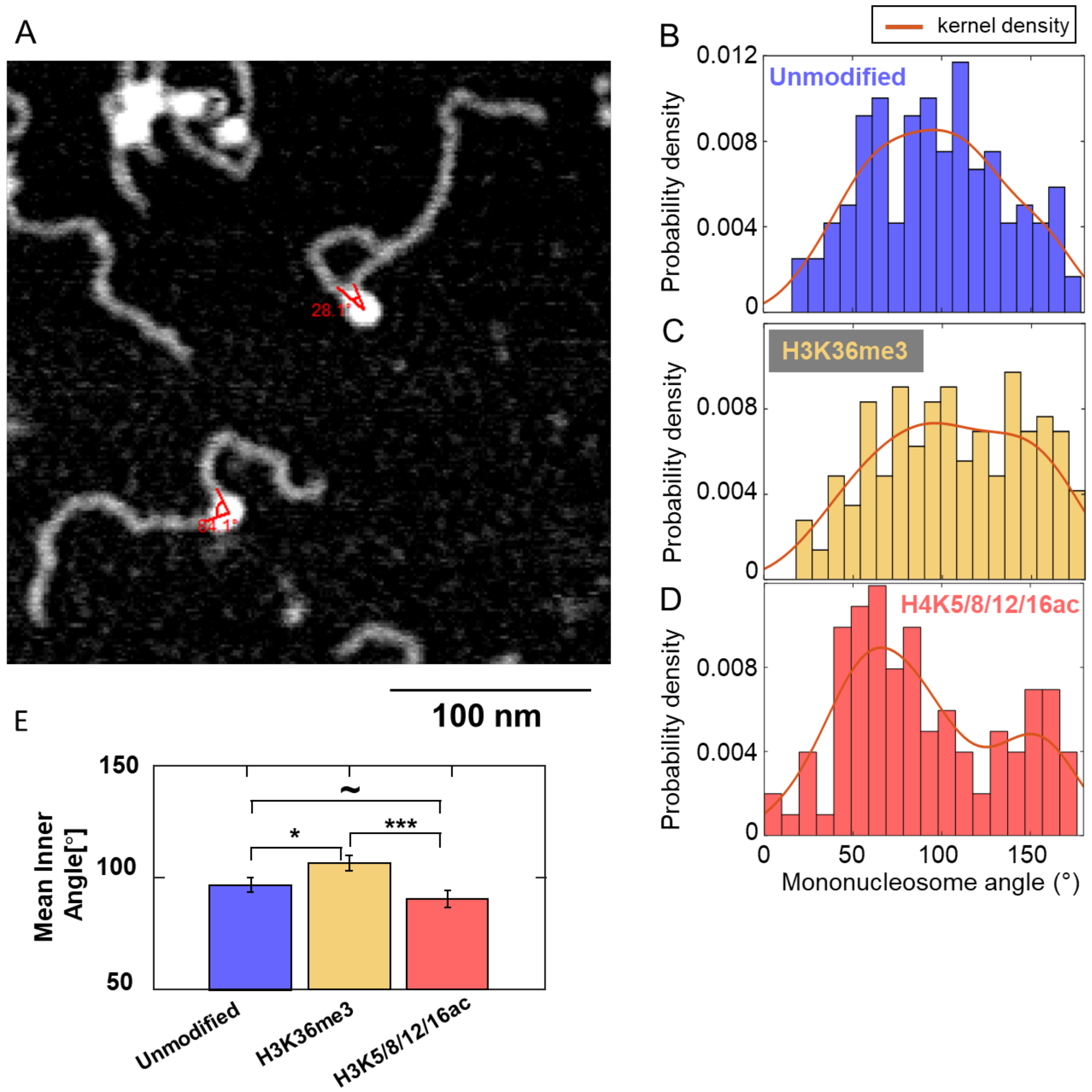


**Supplementary Figure 3. DNA exit angles for mononucleosomes. A)** Example AFM image of mononucleosomes selected from the AFM images of our tri-nucleosomes constructs. The traced DNA exit angles is indicated in the image. **B)** Distribution of exit angles for unmodified nucleosomes (N = 133). **C)** Distribution of exit angles for H3K36me3 nucleosomes (N = 160). **D)** Distribution of exit angles for H4K5/8/12/16ac nucleosomes (N = 120). **E)** Mean exit angles for the three different mononucleosome populations. Error bars indicate the standard error of the mean. The stars indicate significance based on two-tail two-sample *t*-tests: ~ not significant, *p < 0.05, **p<0.01, ***p<0.001.

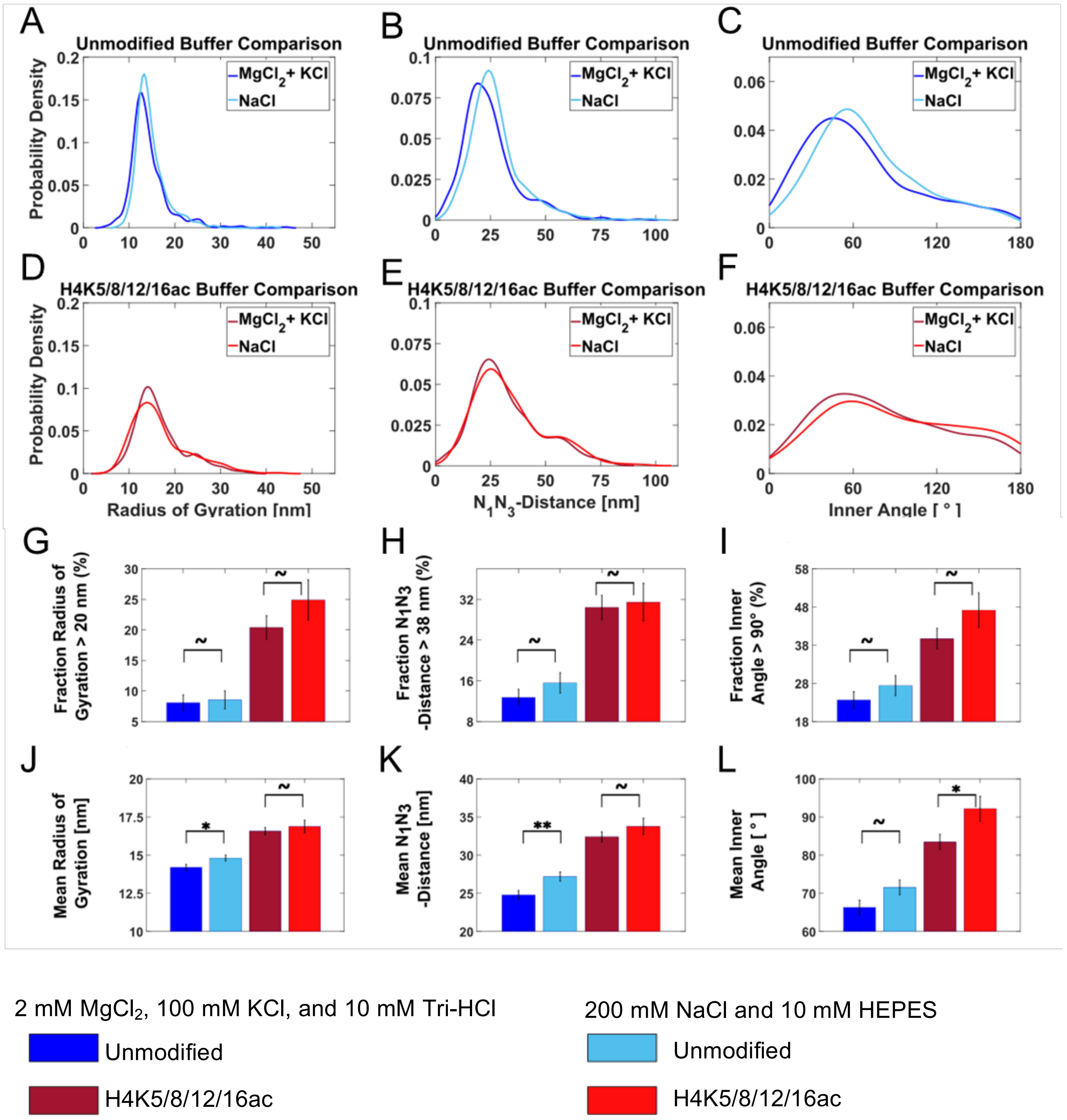


**Supplementary Figure 4. Influence of $Mg^{2+}$ on tri-nucleosome conformations probed by AFM imaging.** Probability distributions as kernel density estimates for radii of gyration, $N_1N_3$-distances, and inner angles for two different salt conditions for unmodified and H4K5/8/12/16ac nucleosomes obtained from AFM imaging. (A-C) Unmodified (D-F) H4K5/8/12/16ac. **(A-F)** show the kernel density estimates of the unmodified nucleosome with different types of salt co-plotted for each parameter. **(G-L)** show analysis of distributions from panels (A-F). ~ not significant, $*p < 0.05$, $**p<0.01$, based on two-tail two-sample proportion-test (G-I) and two-tail two-sample *t*-tests (J-L).

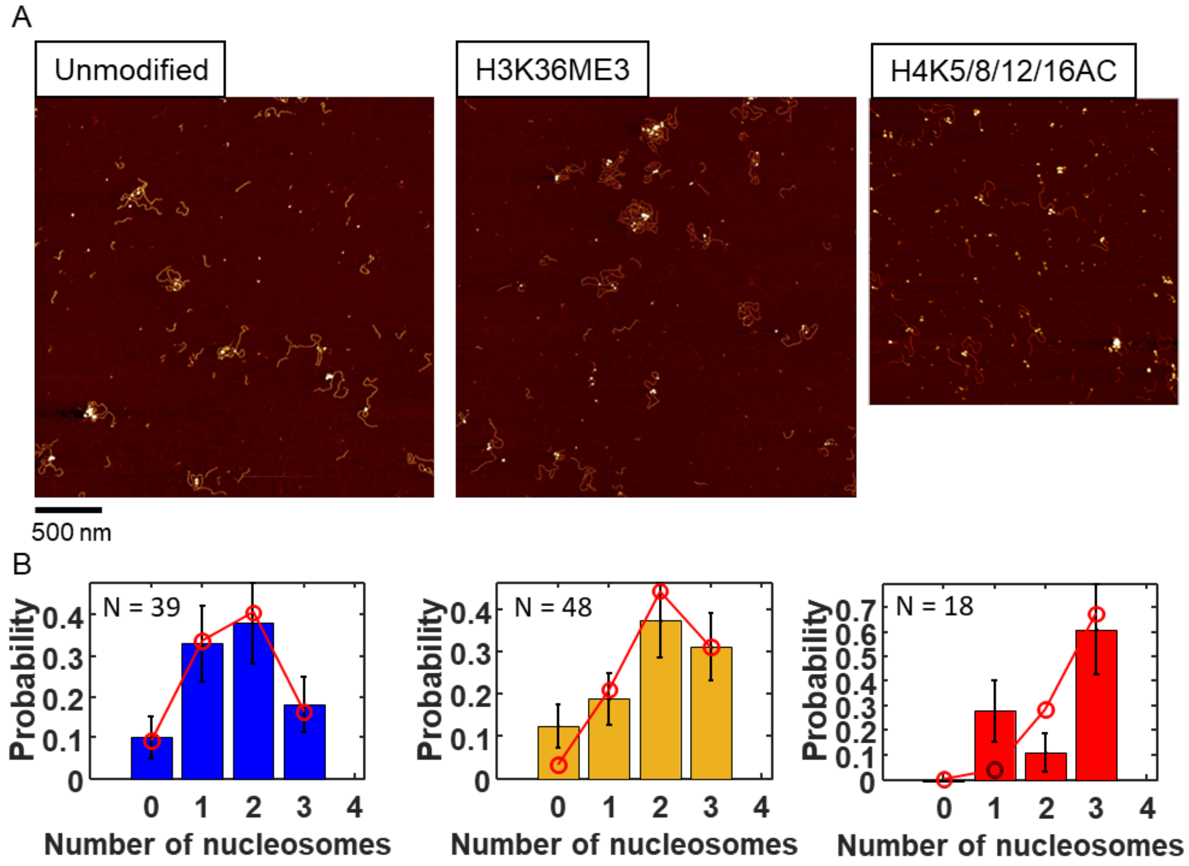


**Supplementary Figure 5. AFM image shows the heterogeneity of nucleosome population. A)** AFM images of unmodified, H3K36me3, and H4K5/8/12/16ac nucleosomes reconstituted on 2823 bp DNA for the MT experiments (left to right). **B)** Histogram of the number of variant nucleosomes assembled on the DNA constructs obtained from AFM images (N, DNA molecules; error bars are from counting statistics). Red points are the best fit of a binomial distribution with fitted assembly probability P = 0.55, 0.68, and 0.88 for unmodified, H3K36me3, and H4K5/8/12/16ac nucleosome reconstitution.

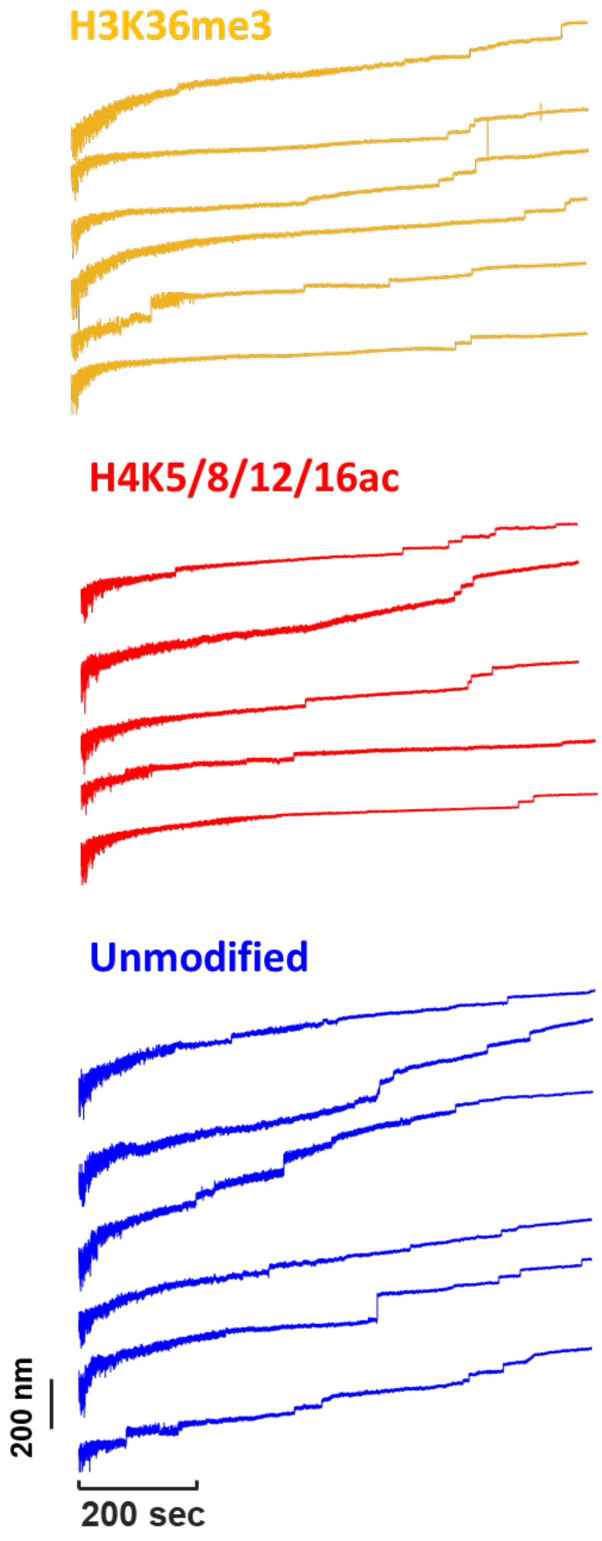


**Supplementary Figure 6. Extension time traces in magnetic tweezers.** Examples of extension time traces under applied force from 0.5 to 30 pN for different variant nucleosomes (Unmodified, blue lines; H3K36me3, orange lines; H4K5/8/12/16ac, red lines).

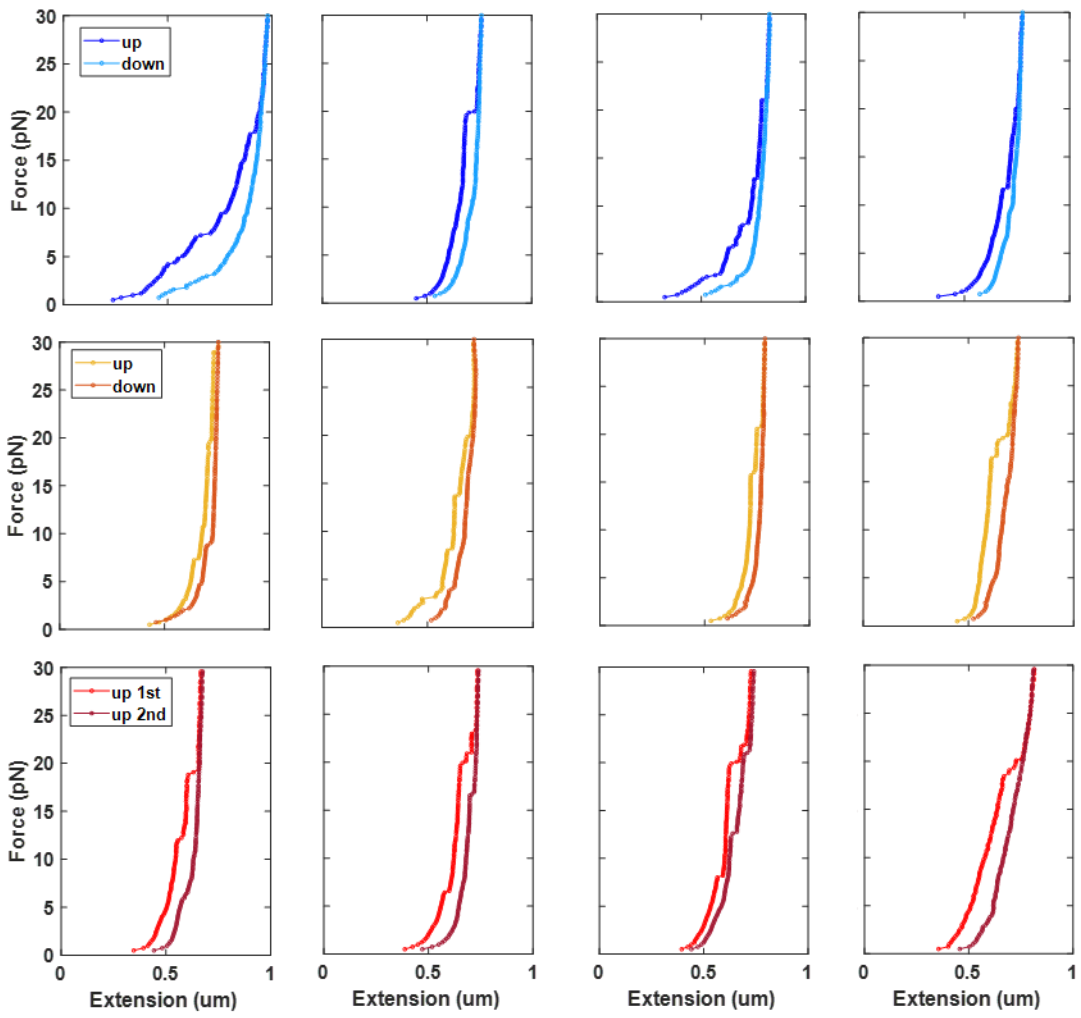


**Supplementary Figure 7. Repeated force-extension cycles for different nucleosome variants.** Examples of force-extension curves for unmodified (top row), H3K36me3 (middle row), and H4K5/8/12/16ac (bottom row) nucleosome constructs. Curves show the first stretch cycle from low to high forces (0.5 – 30 pN; "up") and either the first release cycle (i.e. returning from high force down to low force; "down") or, for the H4K5/8/12/16ac condition, the second stretch cycle ("up 2nd").
